# Obscured Complexity: How External Cycles Simplify the Dynamics of the Endogenous Circadian Oscillator–take the time series of body temperature records as an example

**DOI:** 10.1101/2024.05.09.593452

**Authors:** Feng Lin

## Abstract

**Background:** Understanding circadian rhythms is crucial in various fields of biological research, as they play a fundamental role in the regulation of diverse biological processes, ranging from gene expression to physiological functions.

**Objective:** This study aims to explore the complexity of circadian rhythm signals from a biological system. Without the permission of using experimental data, the mathematical model is utilized to simulate the intricate dynamics of the body temperature’s circadian rhythms and investigate the impact of parameter variation on system behavior.

**Methods:** The Duffing equation is constructed as the mathematical model for simulating circadian rhythms. A thorough discussion justifies the selection of the Duffing equation and establishes the proper parameter range, ensuring chaotic behavior in the system. Four different values of the driving force parameter *γ* (0.32, 0.33, 0.34, and 0.35) are chosen to represent specific cases. Fourier analysis is employed to analyze the simulation data, revealing the frequency components present in the circadian rhythm signals. Entropy analysis along the Poincare sections is utilized to measure the system’s behavior and aggregation of points.

**Results:** The simulations exhibit distinct characteristics in terms of plain visualization, Fourier analysis, and entropy analysis along the Poincare sections. Under normal work sleep conditions (*γ* = 0.35), the system demonstrates specific resetting at particular times within a total period. In shift work (*γ* = 0.34) conditions, some of the resetting behaviour diminishes and the initial phase of the time changes. In longterm constant temperature (*γ* = 0:33) conditions, resembles that of normal work sleep conditions, with a noticeable reset at the beginning of the period. When all external driving forces are eliminated (*γ* = 0:32), the system undergoes multiple resets within a given period. In such circumstances, the biological clock experiences more frequent resets to adapt to the independent operations of each subsystem. Without relying on external environmental cues for regulation, the biological clock relies on frequent resetting to maintain the stability and coordination of the entire system.

**Conclusion:** The simulations reveals variations in resetting behavior and the importance of frequent resets in the absence of external cues. The complexity arising from chaos allows the biological system to adapt and adjust to the intricacies of the external environment. The endogenous clock within the system, despite its inherent complexity, can dynamically optimize its entrainment with external cycles. However, the full complexity of the endogenous clock may be concealed within the system and not readily observable. These findings contribute to a better understanding of the complex dynamics of circadian rhythms. Future research should aim to validate these results through comparisons with experimental data.

## 1 Introduction

Understanding circadian rhythms is crucial in various fields of biological research, as they play a fundamental role in the regulation of diverse biological processes, ranging from gene expression to physiological functions. Researchers have employed a wide range of approaches to investigate circadian rhythms, exploring their intricate dynamics across different levels of biological organization [1–8].

Studies have explored circadian rhythms in various physiological and behavioral phenomena. To study circadian rhythms, researchers employ two main approaches: top-down and bottom-up. The top-down approach involves constructing mathematical models to describe the cyclic changes observed in biological systems. For instance, Kronauer (1999) developed limit cycle models to simulate the oscillatory nature of circadian rhythms [9–11]. Researchers have also utilized Poincare map methods to analyze these non-linear differential equation models, as demonstrated by Diekman and Bose (2016) in their construction of entrainment maps for various circadian models [12–15]. However, challenges remain in developing entrainment maps for arbitrary circadian systems [16]. On the other hand, the bottom-up approach utilizes data mining methods to uncover patterns and information from experimental data. Commonly used techniques include autocorrelation, Fourier analysis, cosinor analysis, and wavelet-based techniques [17–26]. These data processing methods allow researchers to uncover intricate patterns and gain insights into circadian rhythms.

For example, Papavasiliou (1995) employed nonlinear dynamics analysis to study prolactin diurnal secretion, revealing low-dimensional chaotic dynamics. This implies that using a limit cycle model to describe the biological system may not be appropriate. However, there is no further report on the construction of an equation to describe the time series. In recent decades, studies have shown that in caves, the estimated rest-activity cycle period can be longer than the actual rest-activity cycle period, which means that people may feel like they are sleeping for longer than they actually are [27]. Furthermore, research has demonstrated the influence of environmental factors, such as natural temperature changes, in regulating sleep cycles [28]. These findings suggest the need for more complex mathematical models to accurately describe the biological system.

If experimental data were available, we could combine data mining techniques with mathematical modeling to establish a more accurate and quantitative model. Unfortunately, due to the unavailability or limitations of experimental data in certain cases, we may have to rely on qualitative reports to develop a mathematical model.

In the following sections, we will present our proposed qualitative mathematical model and demonstrate how it can explain various observed biological rhythmic phenomena. It is not feasible to construct mathematical models for all biological rhythms; therefore, we have chosen body temperature as a modeling target for the purpose of illustration. Body temperature is a well-known example of a biological rhythm that exhibits circadian patterns. By focusing on body temperature, we aim to develop a qualitative mathematical model that captures the underlying mechanisms driving these rhythmic fluctuations. While our model may not provide precise quantitative predictions without specific experimental data, it offers valuable insights into the dynamics and regulation of body temperature rhythms.

This paper is organized as follows: In Section 2, we employ the Duffing equation as our mathematical model and explore its properties, focusing on the stability analysis of the numerical results. Fourier analysis is used to identify the dominant periods in the system. Instead of studying individual points, we analyze the distribution of points on each Poincare section and quantify their dispersion or aggregation using entropy measures. Section 3 presents detailed results, including simulations of body temperature time series and corresponding entropy along Poincare sections under different scenarios. In Section 4, we analyze and discuss the results, addressing the research questions raised earlier.

## 2 Materials and Methods

### 2.1 Construction of the Duffing equation as a Mathematical Model

#### 2.1.1 Clues and justification

According to numerous studies on circadian rhythms, the suprachiasmatic nucleus (SCN) is widely recognized as a crucial “zeitgeber” for these rhythms [1–4, 29–32]. It serves as the central pacemaker that synchronizes and regulates various biological processes, including the circadian rhythm of body temperature. The SCN acts as the master clock, orchestrating the timing and coordination of physiological and behavioral activities throughout a 24-hour cycle. In the context of body temperature, the SCN plays a crucial role in setting the pace of circadian rhythms. It receives input from environmental cues, such as light exposure, and uses this information to entrain and adjust the timing of body temperature fluctuations. From a mathematical perspective, the regulation of body temperature can be conceptualized as a forced oscillation.

According to a study by Papavasiliou (1995), biological systems exhibit low-dimensional chaotic characteristics [6]. Furthermore, a 2019 report suggests that in caves, individuals may perceive their sleep duration as longer than it actually is, indicating that the absence of a periodic light-dark pacemaker can disrupt circadian rhythms [27]. Another report from 2015 highlights the crucial role of temperature in regulating sleep cycles [28]. These clues suggest that the biological system is far more complex than a simple limit cycle.

Though the experimental data are not allowed to use, the observed results can help us to improve the model. It has been observed that the average temperature under proper work-sleep conditions is slightly higher compared to shift work conditions. Additionally, the amplitude of the fluctuation part of body temperature is larger. The elimination of environmental diurnal temperature changes, can impact the circadian rhythms of body temperature. If all of the external conditions such as temperature changes and light-dark cycles are all been eliminated, the body’s internal biological clock becomes the sole periodic driving force.

#### 2.1.2 Duffing equation

The information provided indeed suggests that circadian rhythms exhibit periodic behavior, and while a simple harmonic oscillator can provide a rough description, it may not fully capture the complexity of the system. Biologists propose that a regulator within the body acts as a forced oscillator, indicating that the oscillation is influenced by external factors and is more intricate than a simple harmonic oscillator. The stability of the oscillator suggests that a simple limit cycle can be appropriate. However the low-dimensional chaotic characteristics of the circadian rhythms suggests that limit cycle may not be appropriate, and a more complex oscillatory behavior is likely involved. Considering these clues, the Duffing equation emerges as a suitable candidate. The Duffing equation is a nonlinear differential equation that describes a forced oscillator with different nonlinear terms, allowing for a more complex oscillatory behavior. This equation can capture the interplay between external forcing, internal regulation, and the nonlinear dynamics exhibited by circadian rhythms. While the Duffing equation is a promising candidate [33], it is important to note that there may be other suitable models that can capture the complexity of circadian rhythms. However, the Duffing equation is a well-studied and understood model that provides valuable insights into the properties and behaviors of oscillatory systems. The Duffing equation can be expressed as:

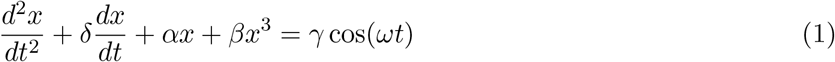

in which *x* represents the system’s displacement. *t* represents time. *δ* controls the amount of damping. *α* controls the linear stiffness, and *α <* 0. *β* controls the amount of non-linearity in the restoring force. When *β* = 0, the Duffing equation returns to a damped and driven harmonic oscillation. *γ* is the periodic driving force. *ω* is the angular frequency of the periodic driving force.

Often, angular frequency of the periodic driving force *ω* is related to external factors such as diurnal temperature and light changes and internal biological clock. The driving force parameter *γ* is influenced by the periodic changes associated with diurnal temperature and light variations as well as the internal biological clock. It is hypothesized that the value of the driving force parameter *γ* is influenced by various periodic changes in the external environment, including diurnal variations in light exposure, temperature fluctuations, and the internal biological clock. The presence of multiple periodic factors leads to an increase in the value of *γ*. Furthermore, if these periodic factors align in phase, it results in a further amplification of *γ*.

### 2.2 Parameter Range Selection of Duffing equation

The model described by Equation (1) exhibits a high level of complexity. When the parameter *γ* = 0, Equation (1) becomes an autonomous system. The equilibrium points are given by 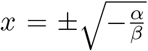. When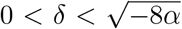, these equilibrium points correspond to stable spirals, while for 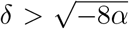, they become stable nodes. Since our focus is on finding periodic solutions, we need to search for shifted 2*π/ω*-periodic solutions in the form described by Jordan and Smith (2007) [34]:

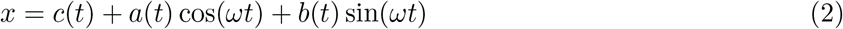

The harmonic balance method is based on the assumption that the amplitudes *a*(*t*) and *b*(*t*) vary slowly compared to cos(*ωt*) or sin(*ωt*) in Equation (1). As a result, their second derivatives can be neglected. Additionally, higher-order harmonics beyond the first will be disregarded. By substituting the approximate solution described in Formula (2) into Equation (1), we can determine the conditions that the undetermined functions *a*(*t*), *b*(*t*), and *c*(*t*) must satisfy:

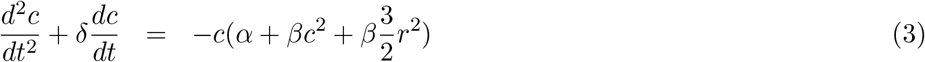

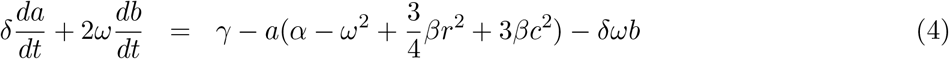

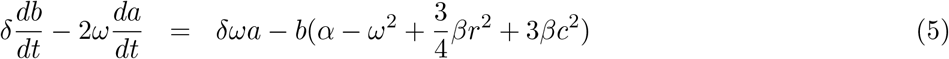

with *r*^2^ = *a*^2^ + *b*^2^. For simplicity, we have omitted the explicit dependence on time *t* in Equations (3)-(5). Although the variables *a* and *b* are actually functions of time, denoted as *a*(*t*) and *b*(*t*), we use the simpler notation *a* and *b* to reduce the complexity of the notation. At steady state condition:

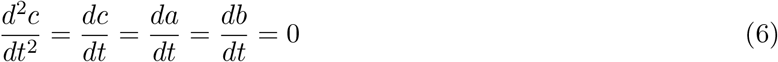

Then

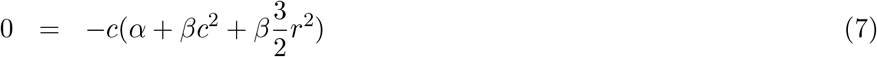

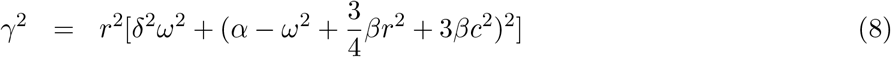

When *c* = 0, the parameters should satisfy

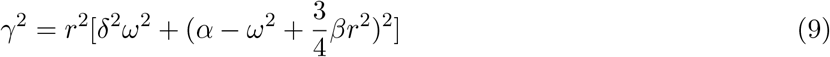

When 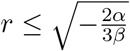 and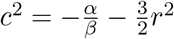, the parameters should satisfy

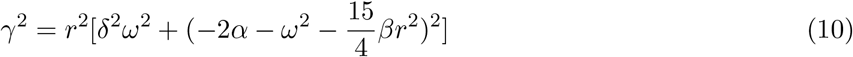

By setting the parameters *α, β, δ*, and *ω* to specific values, such as those mentioned in reference [34], chapter 13 (*α* = *−*1, *β* = 1, *δ* = 0.3, *ω* = 1.2), we can plot curves on the *r − γ* plane. The parameter *γ* plays a crucial role in determining whether the system exhibits periodic or chaotic behavior. Utilizing numerical calculations, we find the following characteristics based on the chosen parameter values:

1. For *γ* ∈ (0, 0.27), stable 2*π/ω*-periodic solutions exist within the system.
2. When *γ* ∈ (0.3, 0.36), the solution displays bounded behavior but does not exhibit periodicity. The Poincare map, used to analyze the system’s behavior, reveals bounded dynamics without obvious repeating patterns. This indicates the presence of chaos within the system.

Additionally, it’s worth noting that the parameters *α, β, δ*, and *ω* can be chosen with different values. However, determining the appropriate values would require additional calculations. For the sake of convenience and to maintain consistency with the reference, we have utilized the parameter values provided by the reference.

In line with our hypothesis, the system exhibits high complexity, and we suggest setting the parameter *γ* within the interval of *γ >* 0.3.

### 2.3 Simulation of time series of body temperature

Given that our model, described by Equation (1), can exhibit either periodic or chaotic characteristics according to the combining of parameters, its solution can be applied to simulate body temperature fluctuations. Typically, the average body temperature ranges from 36 to 37 degrees Celsius, with fluctuations occurring around this average. The numerical solution of Equation (1) *x*(*t*) obtained using the Runge-Kutta algorithm, captures the temperature fluctuations observed within each period.

By setting the average body temperature to be 36 degrees Celsius and considering that the numerical solution is always positive, the body temperature *T* (*t*) varying over time can be expressed as

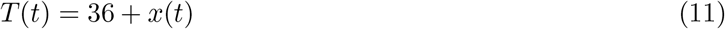

In this equation, *x*(*t*) represents the numerical solution obtained from the model. The angular frequency of the periodic driving force in Equation (1) is denoted as *ω*, and the driving period *τ*_*d*_ satisfies *ωτ*_*d*_ ∈ {1, 2, 3, …}.

### 2.4 Fourier analysis of the time series

The time series data is obtained through numerical solutions of the Duffing equation, as defined by Equation (1), and is represented by Formula (11). To investigate the frequency characteristics of the time series in the frequency domain, we utilize the fft module from the scipy library to perform Fourier analysis. This analysis enables us to identify the dominant frequencies present within the system.

By adjusting the value of *γ* in the Duffing equation, we can simulate different scenarios and examine the corresponding frequency components. This allows us to gain insights into the periodicity or presence of specific frequency patterns under different conditions.

Fourier analysis allows us to extract dominant period of the system, including the initial phase and the ratio of the dominant period’s power spectrum to the total power. These insights help us compare the system’s period change under different values of *γ*, assess any variations in the initial phase, and determine the prominence of the dominant period. The ratio of the dominant period’s power spectrum to the total power helps us evaluate the significance of the dominant period within the overall system. A higher ratio indicates a more pronounced and influential dominant period, while a lower ratio suggests that other frequencies or components play a more substantial role in shaping the system’s behavior. While the dominant period’s power spectrum may have a significant contribution to the total power, it is unlikely to approach a value of 1. This is due to the high complexity of the system, which typically distributes its energy among other signals or frequencies. In other words, the system’s energy is not solely concentrated in the dominant period, but is likely shared with other less apparent or noticeable periodic components. And the initial phase provides insights into the starting point or the relative position of the system within its periodic cycle.

### 2.5 Entropy on each Poincare sections

#### 2.5.1 Poincare section

When the period of the time series are determined, we can get several Poincare sections in a period. Poincare mapping is a dimensionality reduction technique to simplify the analysis of the system, since the Fourier analysis has revealed the existence of stable periodicity in the system. A Poincare section is a lower-dimensional subspace of the original continuous dynamical system that preserves the periodic and quasi-periodic orbits of the system. By observing the orbit of the system over many periods and noting the point at which it first returns to the section, we can create a Poincare map that sends the first point to the second. This map defines a discrete dynamical system with a state space that is one dimension smaller than the original continuous system. We assume that the system is governed by a function *f* (.), which controls the periodic mapping of the system. This function ensures that the trajectory of the system will always return to its original position on a fixed Poincare section. In mathematical language, the function *f* (.) acts on the variable such as temperature *T* (*t*) at any given time *t* is:

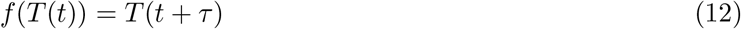

*τ* is the period of the system. For strictly periodic systems, the function *f* (.) controls the periodic mapping of the system and ensures that the trajectory of *T* (*t*) returns to its original position on a fixed Poincare section, i.e. *T* (*t*) = *T* (*t* + *τ*). However, in the presence of external conditions or noise, the location of the returned trajectory may deviate from the original position. *n* times of mapping produces:

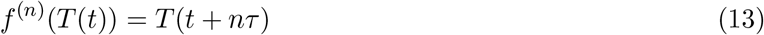

After *n*-times of mapping (*n* = 1, 2, 3, …), *n* points are left on the section. Due to the inherent complexity of biological systems and the presence of noise, the distance between the positions of the point after the *n*th mapping and the initial point will not be zero. However, if the system is stable, it is possible that the distance may converge to a certain region as the number of mappings *n* becomes sufficiently large. To analyze the system using Poincare sections, we observe the orbit in many periods and record the recurrence of the orbit at a definite time in a period [35].

#### 2.5.2 Entropy calculation

According to the points left on the Poincare section, it is possible to figure out the entropy of the distributions on each Poincare sections. In mathematical language,based on the maximum and minimum values of the points, the interval is divided into many intervals at equal intervals *I*_*j*_, *j* = 1, 2, …, *N*. The count of the points in each equal intervals corresponds to the frequency of the temperature *T* in the interval. Divide the count into the total number of points on the thick Poincare section, we get the frequency *p*(*T*). According to the definition of entropy *H*(*T*):

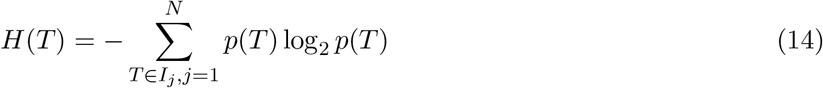

The entropy value reflects the uniformity of the distribution of points across the temperature intervals, with a higher value indicating a more uniform distribution and a lower value indicating a more concentrated distribution in certain intervals. The entropy series of each state provides a measure of the aggregation degree at each Poincare section for that state. By comparing the entropy values of the three stages on each Poincare section, we can assess the degree of accumulation of points over time. Specifically, the more accumulation there is, the lower the entropy value will be.

## 3 Results

### 3.1 Simulation of body temperature in plain visualization

In Figure 1, we present the simulated body temperature obtained from the Duffing equation. As discussed in Section 2, the time series of body temperature records exhibit a high level of complexity, indicating that it is derived from a chaotic system. To explore the influence of the parameter *γ* on the system dynamics, we fix all other parameters and vary *γ* to account for external and internal circadian rhythms. In order to ensure the complexity of our solution, as mentioned in Section 2, we restrict *γ* to the range of (0.3, 0.36). Consequently, we set *γ* to be 0.32, 0.33, 0.34, and 0.35 for our simulations.

**Figure 1:**
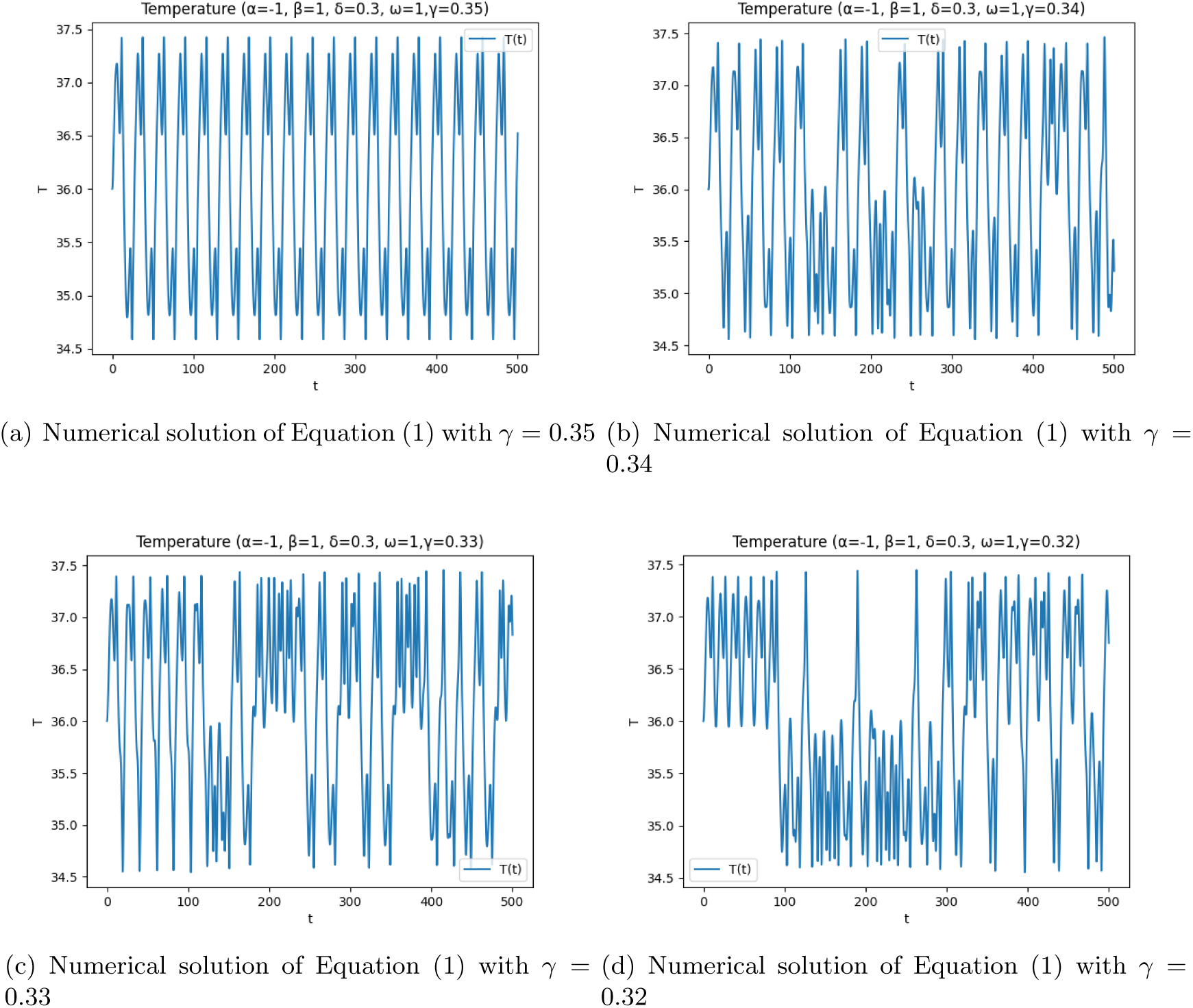
Simulation of diurnal body temperature changes in different cases with parameters *α* = *−*1, *β* = 1, *δ* = 0.3, *ω* = 1.2. Fig 1(a) represents the simulation under normal work sleep conditions. Fig.1(b) represents the simulation corresponding to shift work conditions. Fig.1(c) represents the case of a long-term living in a constant temperature environment. Fig.1(d) represents the case that all of the outside diurnal changes are eliminated. So Fig 1(d) shows that the system exhibits chaotic behaviour.

As discussed in Section 2, the value of *γ* in the Duffing equation is influenced by various periodic changes in the external environment, including diurnal variations in light exposure, temperature fluctuations, and the internal biological clock. The presence of multiple periodic factors contributes to an increase in the value of *γ*. Furthermore, if these periodic factors align in phase, it results in a further amplification of *γ*.

The value of *γ* in the Duffing equation corresponds to different simulated conditions, each with its own implications. When *γ* = 0.35, it represents a simulation under normal work sleep conditions, where the diurnal temperature and light-dark changes align in phase. This aligns with our assumption that a large value of *γ* is expected under this condition. When *γ* = 0.34, it corresponds to a simulation where the sleep-wake cycle is reversed, such as during shift work. In this case, the primary change is the phase of certain factors. For *γ* = 0.33, it simulates a long term living in a constant temperature environment, as supported by the findings in reference [28], where the diurnal temperature changes are eliminated. Finally, when *γ* = 0.32, it represents a scenario where both the temperature and light-dark changes are eliminated, similar to living in a cave for several months. Under this condition, the system exhibits chaotic behavior.

So far, Figure 1(d), which exhibits chaotic behavior, indicates that the circadian rhythms are not apparent in this case. To further analyze its periodicity, we will conduct a Fourier analysis on the time series data.

### 3.2 Fourier analysis of simulations

Table 1 presents the period, power spectrum energy, and initial phase for various values of *γ*. In addition to the cases when *γ* = 0.34 and 0.35, we have also included *γ* = 0.36 and 0.38 for comparison. It is observed that the period remains stable when *γ* is around 0.37, and the initial phase exhibits a consistent ratio of the power spectrum for the most prominent period. However, the ratio of the leading amplitude in the frequency space is not significantly large for *γ* ≥ 0.35, as shown in Table 1. In the analysis of experimental data, it ranges from 0.5 to 0.6, indicating a numerical simulation limitation.

**Table 1:** Period, Ratio of leading power spectrum energy, and Initial Phase.

| $\gamma$ | Period | Ratio | Phase |
| --- | --- | --- | --- |
| 0.38 | 2361 | 0.76 | -1.489 |
| 0.36 | 2361 | 0.812 | -1.512 |
| 0.35 | 2361 | 0.818 | -1.549 |
| 0.34 | 2500 | 0.373 | -3.216 |
| 0.33 | 2174 | 0.147 | -1.404 |
| 0.32 | 50000 | 0.231 | 0.488 |

Interestingly, the multi-period characteristics are more evident when *γ* = 0.34 compared to *γ* = 0.35. The ratio in this case is 0.373, significantly less than 0.818. As is known that the shift work condition alters the phase of work sleep cycles, leading to noticeable changes in the initial phase in Table 1.

For *γ* = 0.33, although the period of circadian rhythm changes is not substantial, the power spectrum of leading period of circadian rhythms is considerably weak. This suggests that long-term living in a constant temperature condition may result in diminished immunological responses.

In the case of *γ* = 0.32, the situation is quite concerning, as the internal biological clock appears to be weaker than anticipated. The period becomes difficult to determine, exhibiting typical characteristics of a chaotic system.

Since there is no specific dimension of time, the period is represented by the count of numbers in our analysis. In our code, the time count is 500, and the step size is 0.01, resulting in a total of 50000 points. The period is determined based on the total number of points and the position of the maximum spectrum. For example, in Table 1, when *γ* = 0.34, the position of the largest spectrum is 20, which implies a period of 2500 counts. This means that when the time and step size are fixed, the period can be compared by the counts.

### 3.3 Entropy distribution along the Poincare sections

In Figure 2, we present the distribution of entropy within a period. As mentioned before, when *γ* = 0.35, it corresponds to a simulation under normal work sleep conditions, where the diurnal temperature and light-dark changes align in phase. In this case, the entropy of the body temperature distribution along each Poincare section will be locally minimized at specific times. This indicates that the external controls help the system to reset, as entropy represents the level of aggregation in the temperature distribution across the Poincare sections.

**Figure 2:**
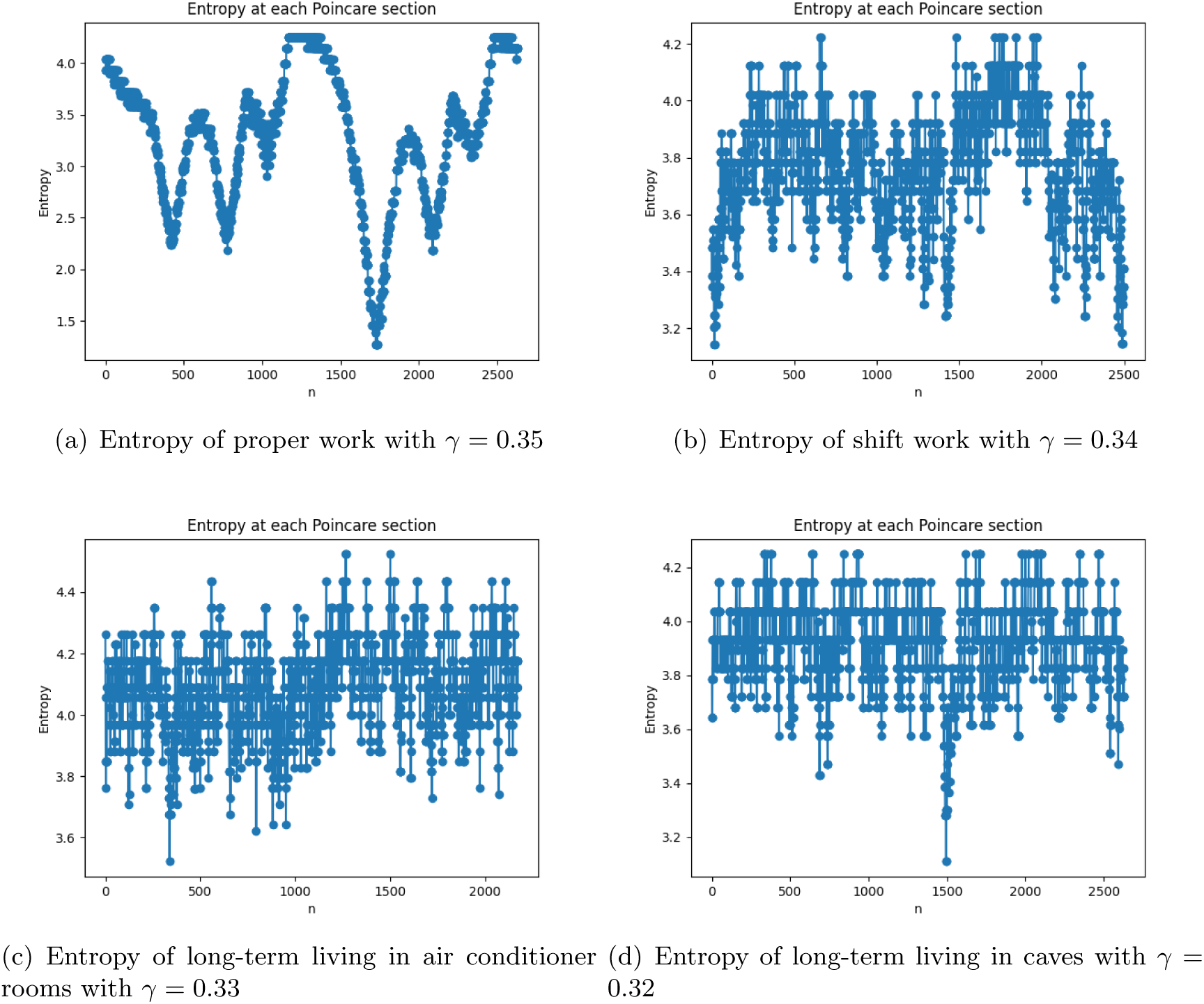
Simulation of entropy distribution in a period with parameters *α* = *−*1, *β* = 1, *δ* = 0.3, *ω* = 1.2. Fig 2(a) represents the simulation under normal work sleep conditions. Fig.2(b) represents the simulation corresponding to shift work conditions. Fig.2(c) represents the case of a long-term living in a constant temperature environment. Fig.2(d) represents the case that all of the outside diurnal changes are eliminated.

When *γ* = 0.34, it represents a simulation where the sleep-wake cycle is reversed, such as during shift work. In this case, the phase of the work sleep cycle does not align with the diurnal temperature and light-dark changes, leading to a discrepancy in the system reset. The reset of the system occurs at noticeably shifted moments compared to normal circadian rhythms, and some reset behaviors may even disappear. These deviations in system reset timing and the loss of certain reset behaviors may be attributed to the disruption of the biological clock caused by the shift in the phase of the sleep-wake cycle. Figure 2(b) illustrates this disruption of order occurring at each end of the period. It indicates a contradiction between the system’s reset according to the diurnal external changes and the shift in the phase of the biological clock.

When *γ* = 0.33, the simulation represents a long-term living scenario in a constant temperature environment, where the diurnal temperature changes are eliminated. Similar to the case of normal work-sleep condition, certain reset behaviors may disappear compared to normal work sleep conditions. Additionally, the reset behavior also resembles that of normal work sleep conditions, with a noticeable reset at the beginning of the period. However, despite this contradiction, the reset phase in this case remains similar to that observed in normal work sleep conditions.

When *γ* = 0.32, it represents the case of living in a cave for several months. In Table 1, there exhibits no obvious period. In order to compare with the above conditions, we have chosen a period length of 2631 counts. In this case, the system exhibits chaotic behavior, and the biological clock undergoes frequent resets. As each cell has its own inherent rhythms of metabolism, the temperature can be seen as the energy dissipated as a result of these metabolic processes. In the absence of external constraints, the lack of a governing system results in each subsystem operating independently. In order to maintain metabolic stability within the overall system, the biological clock has no choice but to constantly reset itself. Under such circumstances, the biological clock undergoes more frequent resets in an attempt to adapt to the independent operations of each subsystem. Without relying on external environmental cues for regulation, the biological clock relies on frequent resetting to maintain the stability and coordination of the entire system.

## 4 Conclusion and discussion

### 4.1 Conclusion

In 1995, Papavasiliou discovered that biological systems might exhibit chaotic behavior. In our study, we employed a schematic mathematical model to simulate the complexity of these systems. Our simulation results align with the analysis of experimental data recording the time series of body temperature, supporting the notion that chaotic behavior is present in biological systems.

Intuitively, one might associate chaotic behavior with sickness or disorder. However, it is important to note that the complexity arising from chaos allows the biological system to adapt and adjust to the intricacies of the external environment. The endogenous clock within the system, despite its inherent complexity, is capable of dynamically optimizing its entrainment with external cycles. It is worth noting that the full complexity of the endogenous clock is often not readily observable or apparent, as it may be concealed within the system.

Though the experimental data are not allowed to use, interested readers have the opportunity to design experiments to verify the predictions presented in this article. By conducting experiments that align with the predictions and hypotheses proposed in this article, researchers can further validate the findings and provide empirical evidence to support the simulation results. This allows for a comprehensive exploration of the topic and reinforces the significance of the study.

By recognizing and studying the underlying complexity and chaotic dynamics of biological systems, we gain a deeper understanding of their adaptability and ability to synchronize with external factors. Further research and analysis can shed light on the intricate mechanisms that govern these systems.

### 4.2 Discussion

The variation of parameters in the Duffing equation significantly influences the system’s behavior, affecting periodicity, bifurcations, and attractor formation. Although this study did not specifically explore the parameter regions associated with periodicity or chaos, it suggests the existence of potentially better parameter values for more accurate simulations.

Furthermore, while the Duffing equation served as a suitable candidate model in this study, there is a possibility that alternative models could provide a better description of the system’s dynamics. To further enhance our understanding of complex biological systems, future research should investigate specific parameter regions linked to periodic or chaotic behavior. Systematic exploration and optimization techniques can be employed to identify optimal parameter combinations that yield more realistic and intricate dynamics.

The discussion on the external and internal driving forces in this study was addressed in a general manner. Further exploration of the driving force mechanisms can improve the model and the construction of parameters.

By incorporating these considerations into future studies, we can advance our understanding of biological systems, their underlying dynamics, and their adaptability to external stimuli.

## Author contributions

Feng Lin constructed the framework, developed the analytical methodology, implemented the codes and write the manuscript.

## Data availability

There is nothing to do with data availability.

## Ethics Statement

This study has nothing to do with experiment.

## Statements

All authors have read and approved the final version of the manuscript.

## References

[1] Satoru Masubuchi, Sato Honma, Hiroshi Abe, Kouji Ishizaki, Masakazu Namihira, Masaaki Ikeda, and Ken-ichi Honma. Clock genes outside the suprachiasmatic nucleus involved in manifestation of locomotor activity rhythm in rats. European Journal of Neuroscience, 12(12):4206–4214, 2000.

[2] Joseph C Besharse. Coupling an activated map kinase to circadian clock output. Neuron, 29(1):3–4, 2001.

[3] Margarita L Dubocovich. Melatonin receptors: role on sleep and circadian rhythm regulation. Sleep medicine, 8:34–42, 2007.

[4] Lawrence P Morin. Neuroanatomy of the extended circadian rhythm system. Experimental neurology, 243:4–20, 2013.

[5] JR Sowers. Dopaminergic control of circadian norepinephrine levels in patients with essential hypertension. The Journal of Clinical Endocrinology & Metabolism, 53(6):1133–1137, 1981.

[6] S. Papavasiliou, T. Brue, P. Jaquet, and E. Castanas. Pattern of prolactin diurnal secretion in normal humans: evidence for nonlinear dynamics. Neuroendocrinology, 62 5:444–53, 1995.

[7] Isabelle Jasper, Andreas Häußler, Barbara Baur, Christian Marquardt, and Joachim Hermsdörfer. Circadian variations in the kinematics of handwriting and grip strength. Chronobiology international, 26(3):576–594, 2009.

[8] Greg Murray and Allison Harvey. Circadian rhythms and sleep in bipolar disorder. Bipolar disorders, 12(5):459–472, 2010.

[9] Megan E. Jewett and Richard E. Kronauer. Refinement of limit cycle oscillator model of the effects of light on the human circadian pacemaker. Journal of Theoretical Biology, 192(4):455–465, 1998.

[10] Daniel B. Forger and Richard E. Kronauer. Reconciling mathematical models of biological clocks by averaging on approximate manifolds. SIAM Journal on Applied Mathematics, 62(4):1281–1296, 2002.

[11] K Tsumoto, G Kurosawa, T Yoshinaga, and K Aihara. Modeling light adaptation in circadian clock: Prediction of the response that stabilizes entrainment. PLoS ONE, 6(6):e20880, 2011.

[12] Casey O. Diekman and Amitabha Bose. Entrainment maps: A new tool for understanding properties of circadian oscillator models. Journal of Biological Rhythms, 31(6):598–616, 2016. PMID: 27754956.

[13] Casey O. Diekman and Amitabha Bose. Reentrainment of the circadian pacemaker during jet lag: East-west asymmetry and the effects of north-south travel. Journal of Theoretical Biology, 437:261–285, 2018.

[14] Jiaxiang Zhang, John T. Wen, and Agung Julius. Optimal and feedback control for light-based circadian entrainment. In 52nd IEEE Conference on Decision and Control, pages 2677–2682, 2013.

[15] C. Papatsimpa, Jochem H. Bonarius, and J.P.M.G. Linnartz. Bio-clock-aware office lighting control. In 2020 16th International Conference on Intelligent Environments (IE), pages 108–114, 2020.

[16] Yuxuan (Nelson) Wu. Challenges of constructing entrainment map for arbitrary circadian models. Honors Theses, page Paper 1350, 2022.

[17] W. Nelson, Y.L. Tong, J.K. Lee, and F. Halberg. Methods for cosinor-rhythmometry. Chronobiologia, 6(4):305–323, 1979.

[18] F.K. Lam, Paul W.F. Poon, A.M.S. Poon, F.H.Y. Chan, and B.M. Wu. Multiscale characterization of chronobiological signals based on the discrete wavelet transform. IEEE Transactions on Biomedical Engineering, 47(1):88–95, 2000.

[19] Joel D Levine, Pablo Funes, Harold B Dowse, and Jeffrey C Hall. Signal analysis of behavioral and molecular cycles. BMC Neuroscience, 3(1), 2002.

[20] Roberto Refinetti, Germaine Lissen, and Franz Halberg. Procedures for numerical analysis of circadian rhythms. Biological rhythm research, 38:275–325, 2007.

[21] Harold B Dowse. Chapter 6 analyses for physiological and behavioral rhythmicity. Methods Enzymol, 454:141–174, 2009.

[22] Tanya L. Leise and Mary E. Harrington. Wavelet-based time series analysis of circadian rhythms. Journal of Biological Rhythms, 26(5):454–463, 2011. PMID: 21921299.

[23] David K. Welsh, Andrew L. Cohen, and Tanya L. Leise. Bayesian statistical analysis of circadian oscillations in fibroblasts. Journal of Theoretical Biology, 314(48):182–191, 2012.

[24] Matthew J. Paul Tanya L. Leise, Premananda Indic and William J. Schwartz. Wavelet meets actogram. Journal of Biological Rhythms, 28(1):62–68, 2013. PMID: 23382592.

[25] Tanya L. Leise. Wavelet analysis of circadian and ultradian behavioral rhythms. Journal of Circadian Rhythms, 11(1):5, 2013.

[26] Tadahiro Goda, Jennifer R. Leslie, and Fumika N. Hamada. Design and analysis of temperature preference behavior and its circadian rhythm in ¡em¿drosophila¡/em¿. J Vis Exp, (83).

[27] Lucrezia Zuccarelli, Letizia Galasso, Rachel Turner, Emily JB Coffey, Loredana Bessone, and Giacomo Strapazzon. Human physiology during exposure to the cave environment: a systematic review with implications for aerospace medicine. Frontiers in physiology, 10:442975, 2019.

[28] Gandhi Yetish, Hillard Kaplan, Michael Gurven, Brian Wood, Herman Pontzer, Paul R Manger, Charles Wilson, Ronald McGregor, and Jerome M Siegel. Natural sleep and its seasonal variations in three pre-industrial societies. Current Biology, 25(21):2862–2868, 2015.

[29] M E Jewett, R E Kronauer, and C A Czeisler. Light-induced suppression of endogenous circadian amplitude in humans. Nature, 350(6313):59–62, 1991.

[30] K Kräuchi. How is the circadian rhythm of core body temperature regulated? Clinical Autonomic Research Official Journal of the Clinical Autonomic Research Society, 12(3):147, 2002.

[31] Premananda Indic, Daniel B. Forger, Melissa A. St. Hilaire, Dennis A. Dean II, Emery N. Brown, Richard E. Kronauer, Elizabeth B. Klerman, and Megan E. Jewett. Comparison of amplitude recovery dynamics of two limit cycle oscillator models of the human circadian pacemaker. Chronobiology International, 22(4):613–629, 2005. PMID: 16147894.

[32] M. A. Hofman. Circadian oscillations of neuropeptide expression in the human biological clock. Journal of Comparative Physiology A, 189(4):823–831, 2003.

[33] G Duffing. Forced oscillations with variable natural frequency and their technical relevance. Heft, 41(42):1–134, 1918.

[34] Dominic Jordan and Peter Smith. Nonlinear ordinary differential equations: an introduction for scientists and engineers. OUP Oxford, 2007.

[35] Amir Shahhosseini, Meng-Hsuan Tien, and Kiran D’Souza. Poincare maps: a modern systematic approach toward obtaining effective sections. Nonlinear Dynamics, 111:529–548, 2023.

